# Manipulating attentional priority creates a trade-off between memory and sensory representations in human visual cortex

**DOI:** 10.1101/2024.09.16.613302

**Authors:** Rosanne L. Rademaker, John T. Serences

**Affiliations:** Ernst Strüngmann Institute for Neuroscience in cooperation with the Max Planck Society, Frankfurt, Germany; Department of Psychology, University of California San Diego, La Jolla, California, USA; Neurosciences Graduate Program, University of California San Diego, La Jolla, California, USA

**Keywords:** visual working memory, distraction, visual attention, fMRI, decoding

## Abstract

People often remember visual information over brief delays while actively engaging with ongoing inputs from the surrounding visual environment. Depending on the situation, one might prioritize mnemonic contents (i.e., remembering details of a past event), or preferentially attend sensory inputs (i.e., minding traffic while crossing a street). Previous fMRI work has shown that early visual areas can simultaneously represent both mnemonic and passively viewed visual information. Here we test the limits of such simultaneity by manipulating attention towards visual distractors during a working memory task performed by human subjects undergoing fMRI scanning. Participants remembered the orientation of a target grating while a distractor grating was shown throughout most of the memory delay. Critically, there were several subtle changes in the contrast and the orientation of the distractor, and participants were cued to either ignore the distractor, detect a change in contrast, or detect a change in orientation. Despite visual stimulation being matched in all three conditions, the fidelity of memory representations in early visual cortex was highest when the distractor was ignored, intermediate when participants attended distractor contrast, and lowest when participants attended the orientation of the distractor during the delay. In contrast, the fidelity of distractor representations was lowest when ignoring the distractor, intermediate when attending distractor-contrast, and highest when attending distractor-orientation. These data suggest a trade-off in early sensory representations when engaging top-down feedback to attend both seen and remembered features, and may partially explain memory failures that occur when subjects are distracted by external events.

## Introduction

Interacting with a dynamically changing environment routinely requires the temporary mental storage of relevant information over short periods of time to guide adaptive behavior – a process that is often referred to as working memory. Depending on task demands, priority can be shifted between maintaining internal representations of recent experiences in working memory, and processing new sensory inputs from the outside world. For example, if you’re meeting a date for the first time, you may want to hold an image of their face actively in mind as you enter the coffee shop, as not to accidentally miss them. However, it is also important to selectively attend your visual surroundings, so that you don’t crash into other coffee shop patrons who are moving about in their singular quest for caffeine. Strategically re-prioritizing either memories of recently viewed information, or directing attention to external visual inputs, is critical to successfully navigating everyday situations.

The abilities to selectively attend to and remember sensory stimuli are thought to be mediated by top-down feedback signals. Specifically, feedback from anterior cortical regions such as the prefrontal cortex (PFC) can modulate neural activity in sensory areas such as early visual cortex (EVC). For instance, selectively attending to a relevant visual feature, such as the eye color of your new date, leads to enhanced firing rates in sensory neurons that are tuned to that feature, along with modulations in neural variability and population covariance structure (Martinez-Trujillo and Treue, 2004; David et al., 2008; Jehee et al., 2011; Liu et al., 2011; Rust and Cohen, 2022), and improved visual sensitivity (Carrasco et al., 2004; Wolfe et al., 2011). In addition to supporting prioritization of relevant sensory inputs, top-down modulations of sensory cortices may also support working memory in the absence of sensory inputs. When a relevant stimulus is no longer present in the environment, top-down signals can engage cortical areas specialized for processing sensory inputs to support detailed memories – an idea termed the *sensory recruitment hypothesis* (Serences, 2016; Adam et al., 2022; Krause et al., 2025). For example, auditory cortex can be recruited to remember an exact pitch (Czoschke et al., 2021; Deutsch et al., 2023), holding an object in mind can recruit object-selective inferotemporal (IT) cortex (Miyashita and Chang, 1988; Miller et al., 1991, 1993; Hirabayashi et al., 2013), and the short-term maintenance of simple visual features, such as particular colors or visual orientations, can recruit early visual cortex (Harrison and Tong, 2009; Serences et al., 2009; Yan et al., 2023). Recruitment of early sensory areas could have functional relevance, as it associated with better working memory recall (Ester et al., 2013; Iamshchinina et al., 2021).

Although selective attention and working memory are thought to recruit the same general areas of early sensory cortex (Van Kerkoerle et al., 2017), it is unclear if they operate via similar or distinct mechanisms and the extent to which they compete. The degree of overlap and competition is unclear because selective attention and working memory have often been studied in isolation. For example, fMRI studies focused on working memory typically use long delay periods that are free from other visual inputs and thus do not mimic real-world scenarios where new images are constantly apprehended by the retina and may require your attention (Harrison and Tong, 2009; Serences et al., 2009; Christophel et al., 2012; Ester et al., 2015; Sprague et al., 2016; Christophel et al., 2018; Kwak and Curtis, 2022). Of course, in real life there are many other visual disruptions such as saccadic eye movements, which often go hand-in-hand with attention. Perhaps sensory recruitment is epiphenomenal, arising only in the absence of concurrent sensory inputs or attentional demands (Leavitt et al., 2017; Xu, 2020).

With respect to concurrent visual inputs, prior fMRI studies found that presenting irrelevant but behaviorally distracting stimuli during the delay period of a working memory task interfered with representations of remembered low-level stimulus features in EVC (Bettencourt and Xu, 2016; Lorenc et al., 2018; Rademaker et al., 2019). However, when salient visual distractors failed to impair recall, interference at the level of EVC proved not obligatory, demonstrating that both behavioral performance and EVC representations can be highly resilient to concurrent visual inputs (Rademaker et al., 2019). Thus, evidence for the degree to which EVC can simultaneously encode new sensory inputs and mnemonic information is mixed. It appears that only when people’s memory performance is negatively impacted, implying their attention was drawn away from their memory contents, interference arises at the level of EVC. This suggests that selective attention and working memory may recruit overlapping cortical resources, eliciting a trade-off between these competing top-down demands.

With respect to concurrent attentional demands, there is indeed some evidence for such a trade-off. Performing a concurrent visual search task can disrupt memory representations of high-level objects such as faces in object-selective areas of ventral visual cortex (Kiyonaga et al., 2017). Moreover, in an fMRI study where participants had to remember a spatial position over a 12 second long delay, a single discrimination performed on a 1 second distractor shown midway through the delay caused a brief dip in the fidelity of the spatial working memory representation in EVC (Hallenbeck et al., 2021). During this transient interference, the remembered position was not entirely erased and remained decodable, just to a lesser extent. Given the brief nature of the distractor stimulus, it remains unclear if working memory contents can be stably maintained in the face of more sustained interactions with external stimuli – common in everyday environments. Moreover, everyday distractions can be more or less engaging, and more or less related to the contents of working memory. Might interference effects depend on the degree of overlap between what is held in working memory, and the type of attention is deployed to the external environment?

Here we test the hypothesis that interference depends on how much priority is strategically allocated to sensory versus mnemonic information. We manipulate concurrent processing demands during the memory delay while using fMRI to measure activation patterns in EVC. We show that activation patterns support decoding of both perceived and remembered information, but that there is a trade-off: Ignoring concurrent sensory input preserves recall performance and the integrity of mnemonic representations in visual cortex, whereas attending to concurrent sensory input leads to lower recall performance and disrupts memory-related activation patterns. This disruption scales with the overlap between remembered and attended features. Together, the behavioral and fMRI results suggest that EVC plays a role in maintaining high-fidelity representations in working memory and that disrupting these representations via the concurrent engagement of visual attention interferes with the successful retention of information in working memory.

## Materials and Methods

### Participants

Nine volunteers (7 female) between the ages of 21 and 32 years (SD = 3.67) participated in the fMRI experiment. Participants had varying amounts of experience with fMRI, ranging from scanner-naïve (S03 and S08) to highly experienced (with > 10 hours in the scanner; S04, S05, and S07). Of our nine volunteers, only the first eight are included for further analyses, as data collection for S09 was stopped at the start of the Covid-19 pandemic. Each of the included volunteers participated in a behavioral training in the lab, and 4–5 two hour testing sessions in the fMRI scanner. For the psychophysical experiment, 32 volunteers were recruited (24 female, mean age = 20.86 + 2.24 years) of whom 17 (12 female, mean age = 21.13 + 2.45 years) were included in the final analyses. Of the excluded participants, 6 did not perform well enough during practice and were never tested, 2 dropped out, 5 were not performing on one or both attention tasks, for 1 there were technical issues, and for 1 data collection was halted due to the start of the COVID-19 pandemic. The study was conducted at the University of California, San Diego (UCSD), and approved by the local Institutional Review Board. All participants provided written informed consent, had normal or corrected-to-normal vision, and received course credit or monetary reimbursement for their time ($20 an hour for fMRI, $12,50 an hour for psychophysics).

### Stimuli and procedure: Main fMRI task

All stimuli were projected on a 16 x 21.3 cm screen placed inside the scanner bore, viewed from ∼40 cm through a tilted mirror. Stimuli were generated using Ubuntu 14.04, Matlab 2017b (Natick, MA), and the Psychophysics toolbox (Brainard, 1997; Kleiner et al., 2007). During the main experiment, memory targets were full contrast donut-shaped sinusoidal grating stimuli (0.75° inner and 10.5° outer radius) with smoothed edges (0.5° kernel with sd = 0.26°), spatial frequency of 2 cycles/°, and mean luminance equal to the grey background. Distractor grating stimuli shown during the delay had the same specifications, with the exception that their contrast was 50% Michelson. Target and distractor grating orientations were pseudo-randomly chosen from one of six orientation bins to ensure a roughly uniform sampling of orientation space (**Supp. Fig. 1**). Importantly, the target and distractor orientations were independent and counterbalanced, such that there was no systematic relationship between them across trials. This counterbalancing was done across 9 consecutive runs of the main task. The recall probe consisted of two black line segments (each 9.25° long and 0.04° wide) such that together the segments spanned the same eccentricity from fixation as the donut-shaped grating stimuli. A central black dot (0.4°) aided fixation throughout.

On every trial we randomly chose a spatial phase for the memory target and the distractor grating (both 0–2). Each initial grating stimulus was then toggled back and forth between its original and inverted contrast at 4Hz, without blank gaps in between, for as long as the grating was on the screen. Thus, the memory target (500 ms total duration) cycled through 1 contrast-reversal (i.e., 250 ms per contrast), such that afterimages were minimized. Similarly, distractors (11s total duration) contrast-reversed for 22 cycles.

Each trial of the main experiment (**Fig. 1a**) started with a 1.6s change in the color of the central fixation dot, indicating with 100% validity the attention condition during the delay (i.e., ignore the distractor, attend changes in distractor contrast, attend changes in distractor orientation). Cues could be blue, green, or red. The pairing of cue-colors with attention-conditions was randomized across participants. Following the cue, a memory target was shown for 500 ms and participants remembered its orientation over a 15 second delay. A distractor grating was presented for 11 seconds during the middle portion of the delay. Irrespective of the attention condition, there would be 2–4 changes in the contrast of the distractor (lower or higher), and 2–4 changes in the orientation of the distractor (counterclockwise or clockwise) on every trial, with each such change lasting for 250ms. Changes could start randomly at any timepoint during the distractor presentation (as long as that timepoint was a multiple of 250ms), with the following exceptions: No changes happened within the first and last 500ms of distractor presentation, and two changes of any kind were always separated by at least 750ms. Participants used the left two buttons on the response box to report each relevant change in the distractor throughout the distractor presentation (i.e., as participants were detecting the changes). During contrast attention trials, participants reported a contrast decrease or increase by pressing the left or right button, respectively. During orientation attention trials, participants reported a counterclockwise or clockwise orientation change by pressing the left or right button, respectively. The total number of changes for each feature (contrast and orientation) was counterbalanced across all 3 attention conditions. After the delay, participants used four buttons to rotate a recall probe around fixation, matching the remembered orientation as precisely as possible. The left two buttons rotated the line counterclockwise, while the right two buttons rotated it clockwise. Using the outer- or inner-most buttons would result in faster or slower rotation of the recall probe, respectively. Participants had 3 seconds to respond before being presented with the next memory target 3, 5, or 8 seconds later. Each run of the main experiment consisted of 12 trials, and lasted 5 minutes and 3 seconds. Data for 36 total runs (432 total trials, with 144 trials per condition) were acquired across 4 separate scanning sessions for all participants (except for S07, for whom all data of the main experiment were collected within the span of 3 longer scanning sessions).

**Figure 1.**
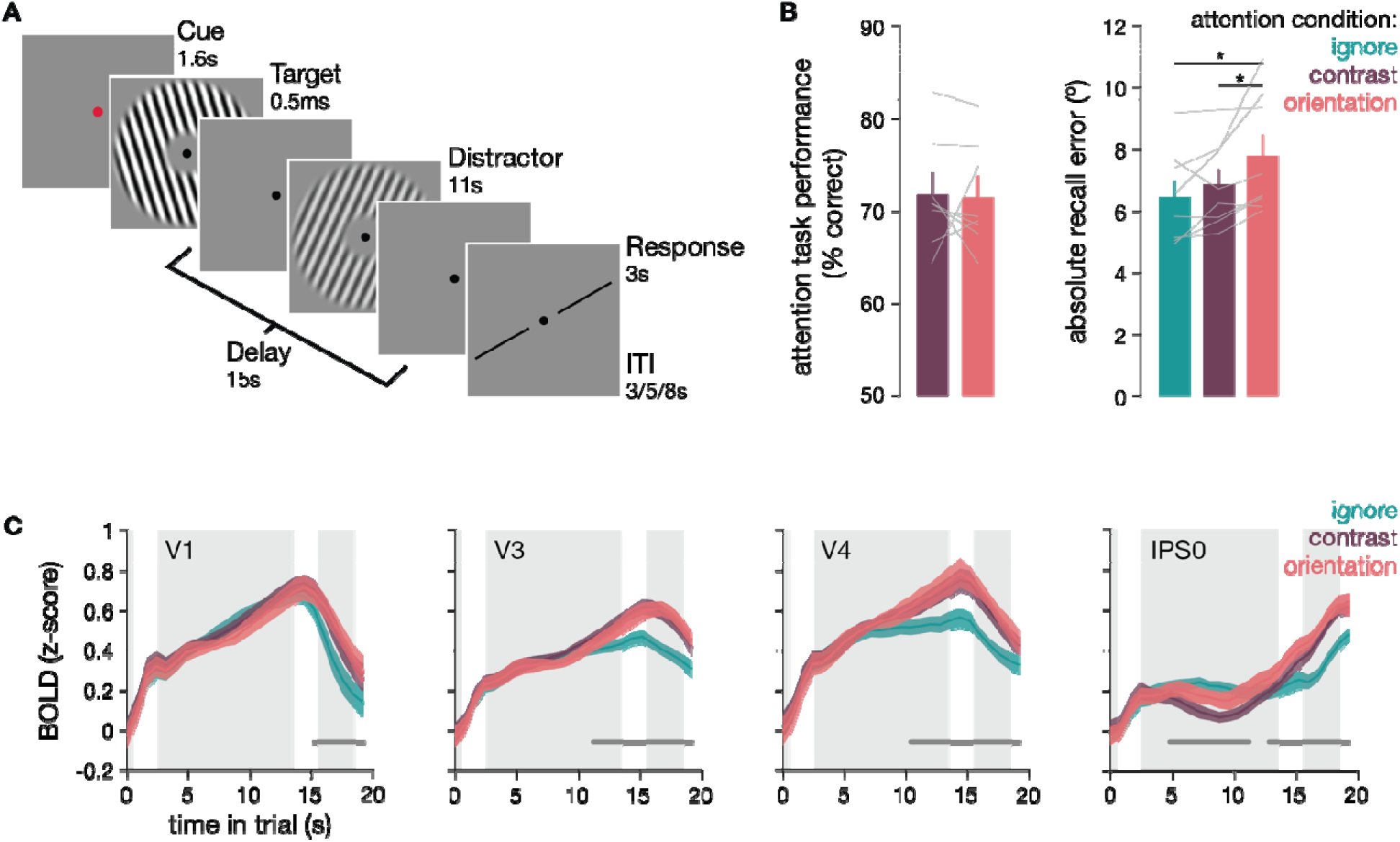
Main task with behavioral and BOLD results. (**a**) Experimental task design. Participant remember the orientation of a brief (0.5s) memory target over a 15s delay, after which they have 3s to rotate a black dial to match the orientation in memory as precisely as possible. On every trial, a distractor grating is shown for 11s during the central-most portion of the delay. On every trial, this distractor i phase-reversing, and has several (2–4) brief (250ms) small changes to its contrast and orientation. Right before the memory target, participants see a 1.6s cue indicating with 100% validity the upcoming attention condition. Specifically, they need to either (1) ignore the distractor grating, (2) attend and report contrast changes, or (3) attend and report orientation changes. When attending the distractor, participants also report the direction of each change (i.e., whether there is an increase / decrease in contrast, or a clockwise / counterclockwise change in orientation). There is an equal number of trials in each condition, and conditions are randomly interleaved. (**b**) Behavioral data recorded while participants were in the scanner shows that performance on the distractor attention task is well-matched (t_(7)_ = 0.157; p = 0.858), and participants perform similarly when detecting contrast (71.85% correct) or orientation (71.56% correct) changes (left panel). Recall of the memory target orientation did differ between conditions (F_(2,14)_ = 5.889, p = 0.002), and was generally worse when participants had to perform a concurrent orientation attention task on the distractor. Bars indicate average performance, and grey lines individual participants. Asterisks indicate significant post-hoc differences between conditions. (**c**) Deconvolved BOLD responses for a few example ROI. Distractors in all 3 attention conditions effectively drove univariate responses in early visual areas, with higher responses when attention was deployed towards the distractor (shown in purple and pink for attention to distractor contrast or orientation, respectively). In IPS0, responses seem more transient, with the strongest response occurring with attention to distractor orientation. Grey background-panels in each subplot indicate the time during which the memory target (0–0.5s, far left panel), the distractor (2.5–13.5s, middle panel), and the response-dial (15.5–18.5s, right most panel) were on the screen. Darker grey lines just above and parallel to the x-axis indicate clusters of consecutive TR’s during which the three attention conditions differ significantly from one another (as calculated with a cluster-based permutation test, see Methods).

**Figure 2.**
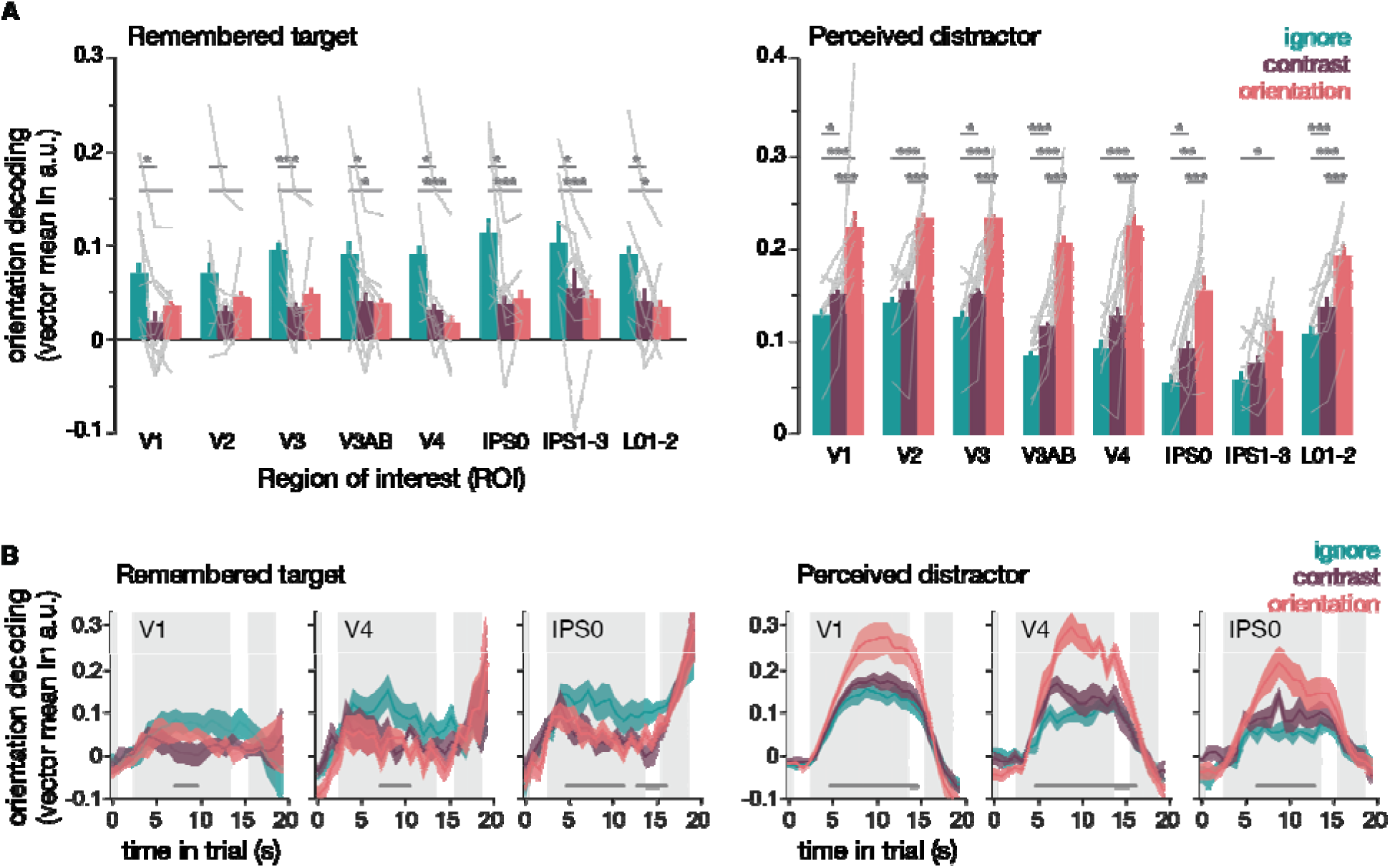
Memory and sensory representations during concurrent top-down demands (**a**) Decoding of the target orientation that is held in memory (left) and of the distractor that was physically presented on the screen (right) during the delay period of the main working memory task. The remembered orientation i better decodable when the distractor is ignored (in teal), compared to when it is attended (in purple and pink). The perceived distractor orientation is least decodable when it is ignored (in teal), better decodable when its contrast is attended (in purple), and best decodable when its orientation is attended (in pink). Bars indicate mean orientation decoding averaged across all participants, while light grey lines indicate individual participants. Dark grey lines and asterisks indicate significant post-hoc differences (*p < 0.05; ** p< 0.01; ***p< 0.001) between attention conditions within each ROI (non-parametric t-tests). Note that for decoding of the remembered target, some significance lines are missing an asterisk indicating significance from within-ROI post-hoc t-tests. Due to the lack of an interaction between condition and ROI, these post-hoc t-tests are not technically warranted, and may be prone to type II errors. Rather, these significance lines indicate the post-hoc tests that compare conditions *across* all ROI, which follow the main effect of attention condition. (**b**) Same decoding as in (**a**) for a few example ROI’s, but shown TR-by-TR throughout the trial of the main working memory task. For decoding time courses of all ROI’s, see **Supp. Fig. 3.** Grey background-panels in each subplot indicate the memory target (0–0.5s, far left panel), distractor (2.5–13.5s, middle panel), and the response (15.5–18.5s, right most panel) periods. Dark grey lines just above and parallel to the x-axis indicate clusters of consecutive TR’s during which the three attention conditions differ significantly from one another (as calculated with a cluster-based permutation test, see Methods).

Before starting the fMRI experiment, participants practiced the main task outside the scanner until they were comfortable using the response buttons to recall the target orientation within the temporally restricted response window; their mean absolute response error for the memory target was <10°; and they were >90% accurate for supra-threshold changes on the distractor attention tasks (both contrast and orientation). Practice took between 5 and 28 blocks of trials, performed across 1–4 separate days.

To equate the difficulty of the two distractor attention tasks, we measured each participant’s performance threshold for both the contrast and orientation attention tasks. This thresholding task was identical to the main experiment, with two exceptions: (1) The “ignore distractor” attention condition was skipped in the interest of time, and (2) a different Δ contrast or orientation was used for every change of the distractor.

Specifically, we used Quest (Watson and Pelli, 1983) to determine the magnitude of change (i.e., contrast increase or decrease; counterclockwise or clockwise orientation deviation) at which participant’s discrimination performance was ∼75% correct. To get an initial threshold estimate, participants completed 1–4 thresholding runs in the lab, prior to scanning. Subsequently, participants also completed 1–2 additional thresholding runs in the scanner prior to each scan session (so while the scanner was still off). This was important for the contrast threshold in particular, since thresholds strongly depend on the specific screen that stimuli are presented on. Once established, these thresholds ensured identical visual inputs irrespective of the attention condition, as the magnitude of contrast and orientation changes was kept constant for each set of 9 consecutive runs of the main experiment. Across participants, the average Δ contrast = 0.116 (SD = 0.075) and the average Δ orientation = 2.074° (SD = 0.547°). As intended, performance on the contrast task (71.85%; SD = 5.848%) and performance on the orientation task (71.56%; SD = 5.718%) did not significantly differ (t_(7)_ = 0.157; p = 0.858; **Fig. 1b**). This means that difficulty was well-equated between the two attention tasks.

### Stimuli and procedure: Localizer tasks

In addition to the main task, we also collected data from an independent sensory localizer task, and an independent memory localizer task. The sensory localizer was used for voxel selection. Both sensory and memory localizer tasks were used for model training purposes (i.e., the analyses shown in **Fig. 3**).

**Figure 3.**
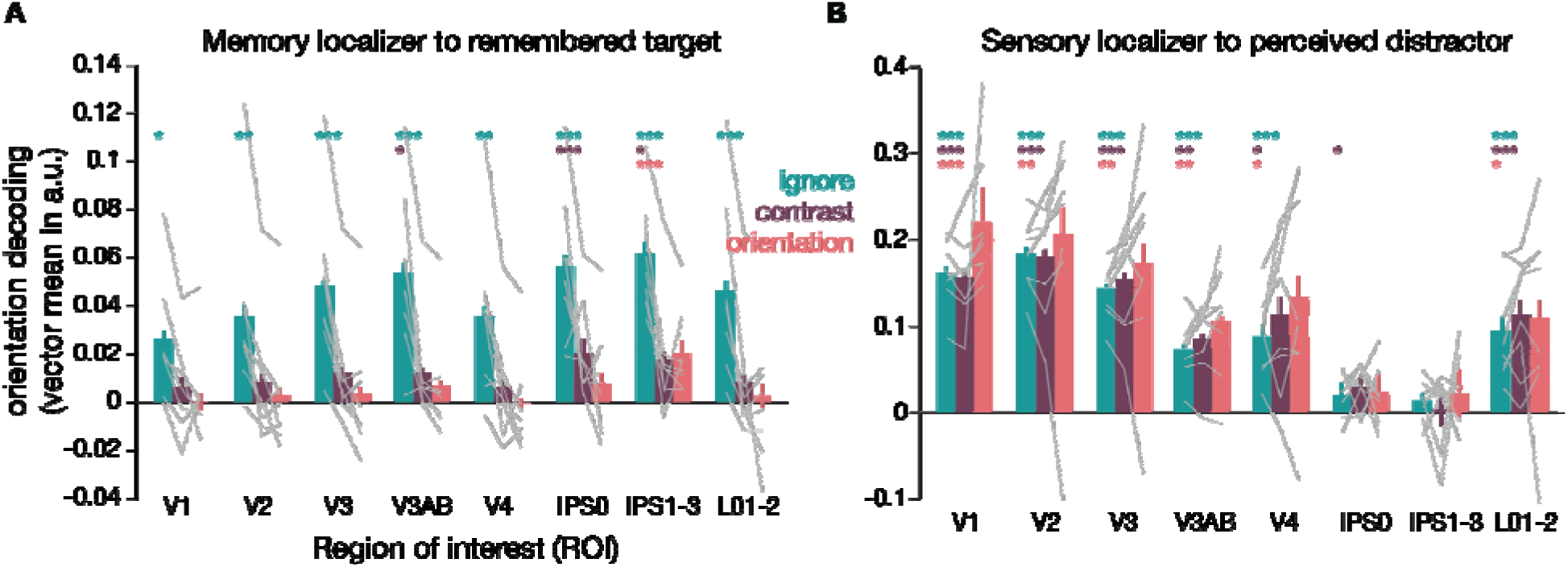
Generalization of memory and sensory formats. (**a**) Decoding of the target orientation that i held in memory, when training a decoder on an independent memory localizer task. The remembered orientation is better represented when the concurrent visual distractor is ignored (teal) than when its contrast (purple) or orientation (pink) are attended. Only in parietal areas (IPS0 and IPS1–3) there i consistent cross-generalization from memory without visual input (i.e., the memory localizer task) to memory with concurrent visual input that is also attended (i.e., the main memory task when the distractor is also attended). (**b**) Decoding of the sensory distractor in the main memory task, when training a decoder on an independent sensory localizer. Response patterns from the sensory localizer task (which used a blob detection task) generalize to responses to the sensory distractor in the main memory task in most ROI, but not in IPS areas. Possibly because of this, no significant differences between the 3 attention conditions are uncovered. Bars indicate mean orientation decoding averaged across all participants, while light grey lines indicate individual participants. Asterisks in matching condition colors indicate significant post-hoc decoding performance (*p < 0.05; ** p< 0.01; ***p< 0.001) compared to chance.

The sensory localizer task consisted of trials showing either a circle-shaped (0.75° radius) or donut-shaped (0.75° inner and 10.5° outer radius) full-contrast grating. On every trial, this grating was shown for 6 seconds (contrast-reversing as in the main experiment), followed by a 3, 5, or 8s inter-trial interval. During grating presentation, a small grey circle was superimposed on the stimulus occasionally (0–3 times per trial) for 250ms. These brief ‘blobs’ could be centered at any distance from fixation between 0.056° and 13.78°, and at any angle relative to fixation (1–360°). Blobs were scaled for cortical magnification such that all blobs stimulated roughly 1mm of cortex (i.e., blobs had radii between 0.18° to 0.75° of visual angle). No blobs were presented during the first 500ms or the last 500ms of a trial, or within 500ms of each other. The blobs functioned to keep participant’s attention directed at the general spatial location occupied by the grating stimuli. Participants were instructed to indicate detection of a blob via a button press. Grating orientation was randomly chosen from one of 9 orientation bins on each trial, and equally often from each bin within a run. Participants completed 36 trials per run (18 circle-shaped grating trials, and 18 donut-shaped grating trials, randomly interleaved), and each run took 7 minutes and 5s. Participants completed between 12–23 total runs of this sensory localizer. These sensory localizer data were obtained in parallel with data collection for another study (Experiment 2 in Rademaker et al., 2019) and have therefore also been used in this previous work.

The memory localizer task was used to estimate voxel responses while participants were remembering an orientation and *not concurrently viewing* any visual inputs (in the main task, memory is always concurrent with visual input from the distractor). During the memory localizer task, participants remembered a briefly presented grating (500 ms; 11.5° radius; full contrast; contrast-reversing as in the main experiment; pseudo random orientation chosen from 1 of 6 bins) over a 12 second blank delay, after which they recalled the orientation by rotating a dial within a 4 s time window. Each trial was preceded by a 1.4s change in the color of the fixation dot (to alert subjects of the upcoming target and indicate the condition), and each trial was followed by a 3, 5, or 8s inter trial interval. For five participants (S01, S04, S05, S06, and S07), these data were collected as part of Experiment 2 in Rademaker et al. (2019), which included trials with and without visual distractors presented during the memory delay. For our current purposes, only the 108 trials with a blank delay period were used (out of the original 324 total trials, collected across 27 total runs). The remaining three participants (S02, S03, and S08) completed 108 total trials of this memory localizer task across a total of 9 runs, and only performed trials without any visual distractors presented during the delay period. Thus, for all participants we have 108 trials with a blank delay period that we use as our memory localizer.

### Stimuli and procedure: Psychophysical experiment

The procedures of the psychophysical experiment largely mirror the procedures of the main fMRI task, with the following exceptions: All stimuli were shown on a 40 x 30 cm luminance calibrated CRT monitor (1600 x 1200 resolution, 100 Hz refresh rate) against a mean grey background (50.3 cd/m^2^), viewed from a distance of 58.5 cm in an otherwise darkened room. Head stability was ensured by using a chin rest. Stimuli were generated under Matlab 2016a and Ubuntu 16.04 LTS. Memory targets and distractors had a 15.5° outer radius (i.e., were slightly larger than in the scanner). Importantly, in this psychophysical experiment a Quest (Watson and Pelli, 1983) staircase was run throughout the entire experiment. This meant that contrast and orientation offsets were adaptively adjusted from one change to the next, ensuring that performance on both attention tasks was equated throughout. Because distractor contrast was 50% Michelson, the biggest possible contrast change was set to 49.5% Michelson, such that the maximum change would show the distractor at near-zero or near-full contrast. Similarly, the biggest possible orientation change was set to 89.5°, because at a >90° offset the difference between two orientations becomes smaller again, due to the circularity of orientation space. At the end of each trial, recall of the remembered orientation was unspeeded, and participants proceeded to the next trial by pressing the space bar. The intervals between trials were shortened by a factor of 1.8, resulting in inter trial intervals of 1.66s, 2.77s, and 4.44s.

As in the main fMRI experiment, counterbalancing was performed across every set of 108 trials. However, instead of presenting participants with a fully counterbalanced set of trials across 9 different runs of 12 trials per run (as in the scanner), for the psychophysical experiment we presented a counterbalanced set of trials across only 3 runs of 36 trials per run. Participants in the psychophysical experiment also performed more trials in total: Instead of 432 total trials (144 trials per condition) as in the scanner, here participants completed a total of 648 trials (216 trials per condition). Participants completed a total of 18 runs over the course of 3-4 days of testing.

### Magnetic resonance imaging

All scans were performed on a General Electric Discovery MR750 3.0T scanner located at the UCSD Keck Center for Functional Magnetic Resonance Imaging (CFMRI). High-resolution (1 mm^3^ isotropic) anatomical images were acquired during a retinotopic mapping session, using an Invivo eight-channel head coil. Functional echo-planar imaging (EPI) data for the current experiment were acquired using a Nova Medical 32-channel head coil (NMSC075-32-3GE-MR750) and the Stanford Simultaneous Multi-Slice EPI sequence (MUX EPI), using nine axial slices per band and a multiband factor of eight (total slices = 72; 2 mm^3^ isotropic; 0 mm gap; matrix = 104 × 104; field of view = 20.8 cm; repetition time/echo time (TR/TE) = 800/35 ms, flip angle = 52°; inplane acceleration = 1). At sequence onset, the initial 16 TRs served as reference images critical to the transformation from k-space to image space. Un-aliasing and image reconstruction procedures were performed on local servers using CNI-based reconstruction code. Forward and reverse phase-encoding directions were used during the acquisition of two short (17 s) ‘topup’ datasets. From these images, susceptibility-induced off-resonance fields were estimated (Andersson et al., 2003) and used to correct signal distortion inherent in EPI sequences using FSL topup (Smith et al., 2004; Jenkinson et al., 2012).

### Preprocessing

All imaging data were preprocessed using software tools developed and distributed by FreeSurfer and FSL (free to download at https://surfer.nmr.mgh. harvard.edu and http://www.fmrib.ox.ac.uk/fsl). Cortical surface gray-white matter volumetric segmentation of the high-resolution anatomical image was performed using the ‘recon-all’ utility in the FreeSurfer analysis suite (Dale et al., 1999). Segmented T1 data were used to generate inflated surfaces on which to draw retinotopic ROIs for use in subsequent analyses. Segmented T1 data were also used for coregistration to the functional data: The first volume of every first functional run of a scanning session was coregistered to the anatomical image. Transformation matrices were generated using FreeSurfer’s manual and boundary-based registration tools (Greve and Fischl, 2009). These matrices were then used to transform each four-dimensional functional volume using FSL FLIRT (Jenkinson and Smith, 2001; Jenkinson et al., 2002), such that all across-session data from each single participant was spatially aligned. Next, motion correction was performed using the FSL tool MCFLIRT (Jenkinson et al., 2002) without spatial smoothing, a final sinc interpolation stage, and 12 degrees of freedom. Slow drifts in the data were removed last, using a high pass filer (1/40 Hz cutoff). No additional spatial smoothing was applied to the data apart from the smoothing inherent to resampling and motion correction. Signal amplitude time-series were normalized via Z-scoring on a voxel-by-voxel and run-by-run basis. Z-scored data were used for all further analyses. Because stimulus onsets were jittered with respect to TR, we aligned the onset of every stimulus to the center of the nearest TR (such that stimuli that started at any time point between -400–400 ms are plotted as time = 0) for every trial (and for every task).

To recover the univariate BOLD time courses for all three attention conditions in the main memory experiment (**Fig. 1c**; **Supp. Fig. 2, top**), and separately for the memory localizer task (**Supp. Fig. 2, bottom**), we estimated the hemodynamic response function for each voxel at each time point of interest (27 TR’s from memory target onset). This was done using a finite impulse response function model (Dale, 1999) consisting of a column marking the onset of each event (memory target onset) with a ‘1’, and then a series of temporally shifted versions of that initial regressor in subsequent columns to model the BOLD response at each subsequent time point. Estimated hemodynamic response functions were then averaged across all voxels in each ROI.

To compute average voxel responses for each trial, to be used for subsequent multivariate analyses, we obtained average activity for the delay period of the main experiment by calculating the mean activation from 5.6–15.2s after target onset (i.e., TR’s 8–20) for every voxel (based on a similar procedure as in Rademaker et al., 2019). For the independent sensory localizer, average responses were calculated over a time window of 3.2–9.6s after grating onset (i.e., TR’s 5–13), and for the independent memory localizer we took the average delay period activation from 5.6–12s after memory target onset (TR’s 8–16).

### Identifying ROI’s

Standard retinotopic mapping procedures (Engel et al., 1994; Swisher et al., 2007) were employed to define eight a priori ROI’s in early visual (V1– V3, V3AB, hV4), parietal (IPS0, IPS1–IPS3), and ventral visual (LO1-2) cortex. Retinotopic mapping data were collected during an independent scanning session, using both meridian (i.e., bowtie-shaped checkerboard stimuli shown at either the horizontal or vertical meridian) and polar angle mapping (a slowly rotating wedge-shaped checkerboard stimulus) to identify voxel’s visual field preferences (described in more detail in Sprague and Serences, 2013). Analyses to obtain retinotopic maps used a set of custom wrappers around existing FreeSurfer and FSL functionality.

To identify voxels that were visually responsive to the part of the visual field where our donut-shaped grating stimuli were shown, a general linear model was applied to data from the sensory localizer using FSL FEAT (FMRI Expert Analysis Tool, v.6.00). Individual localizer runs were analyzed using the brain extraction tool (Smith, 2002) and data prewhitening using FILM (Woolrich et al., 2001). Predicted BOLD responses were generated for each sensory localizer run by convolving the stimulus sequence (of “donut” and “circle” stimuli) with a canonical gamma hemodynamic response function (phase = 0 s, SD = 3 s, lag = 6 s). The temporal derivative was included as an additional regressor to accommodate slight temporal shifts in the waveform to yield better model fits and to increase explained variance. Individual runs were combined using a standard weighted fixed effects model. Voxels that were significantly more activated by the donut compared to the circle stimulus (P = 0.05; false discovery rate corrected) were defined as visually responsive. Only these visually responsive voxels from each ROI were used for subsequent analyses. Exact voxel counts for each participant in each ROI can be found in **Supplementary Table 1**.

### fMRI analyses

For decoding, we used an inverted encoding model (IEM). We chose this approach because it can be used to model continuous feature spaces, in this case orientation, and does not require discrete binning. For our main analyses (**Fig. 2** and **Supp. Fig. 3**) model training and testing was performed using a 4-fold cross-validation procedure. Specifically, each observer participated in a total of 36 runs (with 12 trials per run) of the main memory task, with complete counterbalancing of orientation bins and task conditions per set of 9 runs (a single session). Thus, for model training and testing on each fold, 27 runs (324 trials) served as the training set, and 9 runs (108 trials) were held out as the test set. Each set of 9 runs was held out once. Model training was done on the average pattern activity across delay period TR’s, and across all 3 attention conditions to ensure that we used the same training data throughout. This is different for the results in **Fig. 3**, where we did not use cross-validation but instead trained the model on data from independent localizers.

The first step of the IEM estimates an encoding model by modelling the response in each voxel to 9 raised cosine orientation filters, or “channels”, to characterize the orientation sensitivity profile for each voxel based on training data. The encoding model for a single voxel has the general form:

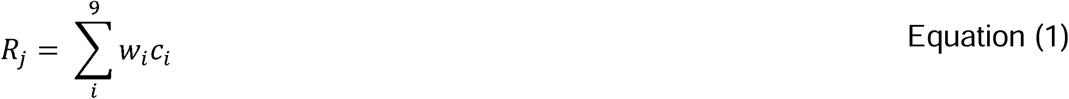

Where *R_j_* is the response *R* of voxel *j*, and *c_i_* is the channel magnitude *c* at the *i*^th^ channel. A voxel’s orientation sensitivity profile is captured by 9 weights *w*, one for each channel. Channels were modeled as:

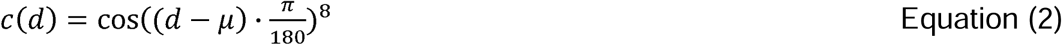

Where *d* is the distance in degrees from the channel center *μ*. Channel centers were spaced 20° apart. Equation 1 can be expressed for matrices as:

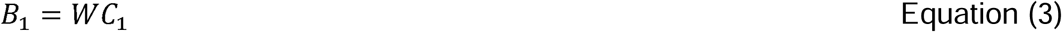

Here, a matrix of observed BOLD responses *B*_1_ (*m* voxels x *n* trials) is related to a matrix of modeled channel responses *C*_1_ (*k* channels x *n* trials) by a weight matrix *W* (*m* voxels x *k* channels). For each trial, *C*_1_ is the pointwise product of a stimulus mask (i.e., “1” at the true stimulus orientation, “0” at all other orientations) by channels. *W* quantifies the sensitivity of each voxel at each idealized orientation channel, and can be computed with least-squares linear regression:

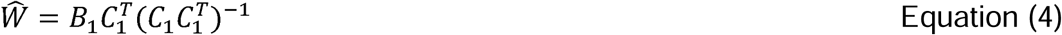

Estimating the sensitivity profiles concludes the first encoding step of the IEM. The second step of the IEM inverts the model, using the estimated sensitivity profiles of all voxels *Ŵ* (*m* voxels x *k* channels) in combination with a test set of novel BOLD response data *B*_2_ (*m* voxels x *n* trials) to estimate the amount of orientation information at each channel *Ĉ*_2_ (*k* channels x *n* trials):

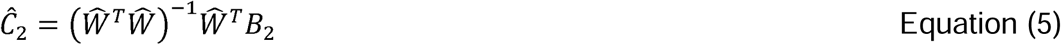

This step uses the Moore-Penrose pseudoinverse of *Ŵ*, and uses the sensitivity profiles across all voxels to jointly estimate channel responses *Ĉ*_2_ for each trial of the test set.

Grating orientations could take any integer value between 1° and 180°. To estimate channel responses *Ĉ*_2_ for each degree in orientation space, both the encoding (Equation 4) and inversion (Equation 5) steps of the IEM were repeated 20 times. On each repeat, the centers of the 9 channels (Equation 2) were shifted by 1°, and we estimated the channel responses *Ĉ*_2_ at those 9 centers, until the entire 180° orientation space was estimated in 1° steps. After generating model-based channel-responses for each trial, all single trial channel-responses were re-centered either on the remembered orientation, or on the orientation of the distractor grating.

Model-based channel-responses were quantified using a “vector mean” fidelity metric derived from trigonometry (Rademaker et al., 2019), and applied to the average of 144 single-trial reconstructions per attention condition (for each condition, participant, and ROI). This vector mean fidelity metric was calculated by convolving the channel response with a cosine, which is equivalent to projecting the channel response at each degree in orientation space onto the center of that space (0°) and taking the mean over all 180 projected vectors. The vector mean reflects if there is modeled information about a remembered or perceived orientation in the channel response profile (it will be zero if not). Note that this metric, by design, gets rid of additive offsets.

### Statistical procedures

To compare participant’s behavioral performance across the 3 attention conditions we used a non-parametric 1-way ANOVA. This was followed up with non-parametric two-sided t-tests to compare performance between each pair of conditions. The same t-test was used to compare performance on the contrast and orientation attention tasks. Specifically, non-parametric tests used 1000 permutations. On each permutation we randomly shuffled the condition labels belonging to the mean performance from each participant, and calculated the appropriate test statistic (an F-value in the case of the ANOVA; a t-value in the case of the t-test) across all participants. Across all 1000 permutations we obtained a null distribution of shuffled test-statistics, against which we compared the test statistic calculated from the intact data to determine the significance level.

To compare decoding performance across the 3 attention conditions and across our 8 ROI’s, we performed a permutation-based 2-way ANOVA, which works similarly to the 1-way ANOVA described above (condition labels are shuffled, and a test statistic is calculated across 1000 permutations). In the case of a significant 2-way interaction between condition and ROI, we also test for differences between the 3 attention conditions *within* each ROI (again using non-parametric t-tests). Note that while technically there was only one such 2-way interaction, namely, for distractor decoding (**Fig. 2a**, right), we also performed these post-hoc tests for memory decoding (**Fig. 2a**, left) for consistency in the figure. To follow up on main effects of attention condition (**Fig. 2a**, **Fig. 3a**), we tested for differences between conditions by taking the average vector mean for each participant across all ROI’s, and then performing pair-wise post-hoc tests between all 3 conditions using a non-parametric t-test as already described above. Finally, to follow up on main effects of ROI, we checked for significant above-chance decoding in each condition and ROI (**Fig. 3**) by performing one-sided non-parametric t-tests against zero.

To determine if univariate BOLD responses or decoding differed between the 3 attention conditions over time (i.e., TR’s within a trial), we used a non-parametric 1-way ANOVA cluster-based permutation test. Specifically, to verify at which timepoints there were significant differences between the 3 attention conditions, we used a threshold of p = 0.05 (F = 2.73) and calculated the observed cluster mass (i.e., sum of F-values for adjacent timepoint that supersede the threshold) for every cluster in every ROI. Then, we generated a null-distribution of cluster mass sizes: Across 1000 permutations we shuffled the condition labels for the data in each ROI, while keeping the temporal structure of the data intact, taking the largest cluster mass on each permutation. We then compared the cluster mass at each *real* cluster in the data against this null-distribution. This cluster-based permutation test is performed for each ROI separately.

For the psychophysical experiment we looked at the circular mean and SD over the entire orientation space (**Supp. Fig. 4 A**, **C**, and **D**), which can be quite noisy at an individual participant level with an orientation space of 180° and only 216 trials per condition (∼1.2 trials per degree). To remedy this, we first used a sliding window of +10° such that the estimate at each degree contained ∼25 trials. Second, we performed bootstrapping to estimate the reliability of these results. Specifically, on each iteration we drew a random sample of participants (with replacement) and calculated the average circular mean or SD at each degree in orientation space. We then calculated the 95% confidence interval (CI) across a total of 1000 bootstraps. This allowed us to evaluate if bias or SD differed significantly in different parts of the orientation space.

### Glitches

We re-measured the screen size and viewing distances in the scanner since the publication of Rademaker et al., 2019, which led to slightly different stimulus sizes reported here for our sensory localizer, compared stimulus sizes reported in reported in Rademaker et al., 2019 (where the same data were used as “second mapping task” in Experiment 2). Partway through data collection, the bulb in the projector burnt out and needed to be replaced. After replacing the bulb, contrast thresholding had to be re-done for affected participants as the new bulb was considerably brighter than the old one. Finally, for S06, we had to abort what was supposed to be the 3^rd^ session after only 3 runs of the main task. This entire session was re-run on another day so that S06 had a complete data set. Data collection for the psychophysics experiment was aborted due to the pandemic (midway through collecting S32) and was never resumed due to the first author moving institutions. Thus, we have fewer participants than was planned a priori.

## Results

First, we wanted to verify that potential neural differences between the contrast and orientation attention conditions in the main working memory task would not be due to differences in task difficulty. Indeed, thanks to a strict staircasing procedure, performance on the contrast (71.85%) and orientation (71.56%) attention tasks did not significantly differ (t_(7)_ = 0.157; p = 0.858; **Fig. 1b**, left panel). Next, we looked if participant’s recall performance during the main working memory task was impacted by concurrent attention to a distractor shown throughout most of the delay (i.e., for 11s). Recall performance differed significantly between the 3 attention conditions (F_(2,14)_ = 5.889, p = 0.014; **Fig. 1b**, right panel), with larger recall errors when participants attended orientation changes in the distractor, compared to when they attended contrast changes (t_(7)_ = 2.623, p = 0.018) or ignored the distractor (t_(7)_=2.53, p = 0.022). Recall errors did not differ significantly between trials during which participants attended contrast changes compared to when they ignored the distractor (t_(7)_=1.497, p = 0.182). Note however that in an independent psychophysics experiment with a larger number of participants (see later in the Results) we *did* observe that recall errors were also larger with contrast attention.

Next, we looked at univariate responses in the main working memory task to see if distractor attention impacts the BOLD time course. We show these univariate time courses for several ROI’s in **Fig. 1c** (for all ROI time courses see **Supp. Fig. 2, top row**). It is clear that the 11s distractor drives a large and sustained BOLD response, especially in early visual areas (and when compared to the memory localizer, where the delay was without visual input, see **Supp. Fig. 2, bottom row**). Towards the end of the trial in particular there appears to be a larger univariate response when distractor attention is required, compared to when the distractor is ignored. Note that in V1 this difference between conditions only emerges 12.8s after the onset of the distractor, meaning that differences in response preparation (memory recall) could also play a role.

Next, we tested if we could decode the orientation of the memory target that was held in working memory, while also concurrently decoding the orientation of the distractor grating that was physically on the screen. Here, we used the average pattern response during the delay to train and test our decoder (an IEM, see Methods). The ability to decode the memory target orientation during the delay differed between attention conditions (main effect: F_(2,14)_ = 10.814; p < 0.001) which was the case for all ROI’s (no condition x ROI interaction: F_(14,98)_ = 1.024; p = 0.439, and no main effect of ROI: F_(7,49)_ = 1.048; p = 0.417)(**Fig. 2a**, **left**). Specifically, the remembered orientation was better decoded when the distractor was ignored, compared to when participants attended changes in distractor contrast (t_(7)_= 3.925; p < 0.001) or orientation (t_(7)_= 3.586; p = 0.012), without a significant difference between the contrast and orientation attention conditions (t_(7)_= 0.212; p = 0.814). These decoding results from visual cortex mirror the differences we see in recall behavior, where performance is also best when the distractor can be ignored.

The ability to decode the orientation of the distractor shown during the delay similarly differed depending on the attention condition (main effect: F_(2,14)_ = 16.08; p < 0.001), but these differences were not the same in all ROI’s (condition x ROI interaction, F_(14,98)_ = 2.155; p = 0.008, and main effect of ROI: F_(7,49)_ = 24.781; p < 0.001)(**Fig. 2a**, **right**). Post-hoc tests in each ROI show that while we observed a difference between attention conditions in every ROI (all F > 4.5; all p < 0.024), all 3 attention conditions differed from one another in some ROI’s but not in others. In general, decoding of the distractor grating was highest when participants had to pay attention to orientation changes – both compared to when participants paid attention to contrast (t_(7)_= 3.34; p < 0.00) and when they ignored the distractor (t_(7)_= 3.34; p < 0.001). Also attention to contrast changes generally improved decoding compared to when the distractor grating was ignored (t_(7)_= 2.961; p = 0.024).

We subsequently evaluated how these differences evolve over time by looking at IEM decoding for each TR of the main working memory trial (**Fig. 2b**; **Supp. Fig. 3**). Despite the increase in noise of single TR data, we nevertheless observed time clusters during which the 3 attention conditions diverged, in line with the results in **Fig. 2a**. Orientation decoding of both the memory target and distractor gratings differed depending on the attention condition, and these differences emerged around 6–8 seconds into the trial, likely reflecting the change in cognitive state at the onset of the distractor grating.

In the previous analysis, we trained our decoder across all 3 conditions of the main working memory task to see how well we could retrieve information about remembered and perceived orientations under different attentional conditions. The benefit of this is that we were not biasing our decoder in favor of any one condition. But there are also downsides to such an approach. For example, the signal-to-noise (SNR) ratio may be worse when we ask participants to perform two competing top-down tasks (i.e., remembering an orientation while also paying attention to concurrent visual input). By training our decoder on such data, we may not be able to uncover all possible differences between the 3 attention conditions. Another notable downside of training and testing a decoder within the same task is that we remain agnostic to the representational format used to remember or process visual inputs. We just know that activation patterns during the delay can predict the remembered orientation, but we do not know if those patterns are similar to memory representations under conditions without visual input. Similarly, we know that activation patterns during the delay (and while the distractor is shown) can predict the perceived distractor orientation, but we do not know if those patterns are similar to sensory driven responses evoked under different circumstances. Thus, we next used data from two independent localizer tasks to see if response patterns generalize from other working memory and perception tasks to the patterns we measured in our main working memory task – when participants were concurrently remembering and perceiving two different orientations.

Do working memory representations measured *without* concurrent visual input (i.e., during a blank delay period) generalize to the memory representations in our main task, where orientation was remembered concurrently with visual distraction on every trial? When we train on data from the independent memory localizer, we can indeed decode the memory target orientation during visual distraction, though this decoding differed between attention conditions (main effect attention condition: F_(2,14)_ = 41.248; p < 0.001), which was true for all ROI’s (no condition x ROI interaction, F_(14,98)_ = 1.683; p = 0.061)(**Fig. 3a**). As before, ignoring the distractor resulted in better decoding of the orientation in memory compared to when distractor contrast (t_(7)_= 6.66; p < 0.001) or orientation (t_(7)_= 6.803; p < 0.001) had to be attended. The remembered orientation was represented marginally better when participants attended contrast as opposed to orientation changes in the distractor (t_(7)_= 2.311; p = 0.054). Decoding performance also differed between ROI’s (main effect ROI: F_(7,49)_ = 9.877; p < 0.001), and was generally higher in IPS. While this may arise from less interesting factors such as SNR differences between ROI (e.g., due to differences in the underlying organization of orientation selectivity), it is noteworthy that such differences were *not* observed when training and testing a decoder within the same task (**Fig. 2a**, **left**). Post-hoc tests further showed consistent above-chance memory decoding when distractor attention was required only in parietal cortex (IPS0 and IPS1–3) (**Fig. 3a**). Thus, the ability to cross-generalize from memory during a blank delay, to memory during a delay with visual distraction, implies a common format for remembering orientation under these different circumstances. Importantly, this common format appears only when the distractor input is ignored, with the exception of parietal cortex, where we see that this overlap in representational format exists irrespective of whether the visual distractor was ignored or attended.

Second, we wanted to know if response patterns generalize from independently measured sensory driven responses, to sensory distractor representations measured under a concurrent working memory load and with different forms of distractor attention. During the sensory localizer, participants performed a behavioral task that was deliberately orthogonal to all 3 of the attention conditions in the main task. Specifically, participants had to detect small gray blobs that could be superimposed anywhere on the sensory grating. This ensured that the localizer stimulus was not ignored, and also that neither contrast nor orientation were attended. The ability to decode the distractor orientation during the delay did not significantly differ between attention conditions (main effect attention condition: F_(2,14)_ = 2.199; p = 0.133) (**Fig. 3b**). The most probable reason for this is the complete lack of decoding in parietal areas IPS0 and IPS1–3 (main effect ROI: F_(7,49)_ = 19.574; p < 0.001). And while there is no significant interaction between attention condition and ROI (F_(14,98)_ = 743; p = 0.741) differences between conditions can be harder to uncover when decoding is absent in some ROI’s (and therefore cannot show any condition differences). It is interesting that, unlike in our main analysis (**Fig. 2a**, right), there is no information about the distractor in IPS. This lack of cross-generalization between the sensory localizer and the sensory distractor in IPS is consistent with a change in the representational format between the different tasks that are performed while sensory inputs are presented. Namely, blob detection without memory load in the sensory localizer, versus an extent of attention with simultaneous memory deployment in the main memory task.

In the scanner, we observed that attending to the orientation of the distractor led to higher behavioral recall errors compared to the attend contrast and the ignore conditions, while in the attend contrast condition recall errors were only numerically (but not significantly) higher than in the ignore condition (**Fig. 1B**). To replicate and further explore this effect, we ran a follow-up behavioral study with a larger sample size to better understand the interactions between attention to the distractor and memory recall errors. We ran the main working memory task (**Fig. 1A**) as a separate psychophysical experiment and we looked at memory recall errors in 17 participants who performed a total of 648 trials (216 per condition) each. To equate performance on the contrast and orientation attention tasks we continuously staircased performance throughout all trials (**Fig. 4A**), which resulted in an average contrast = 0.198 (SD = 0.051) and an average orientation = 3.513° (SD = 2°). As intended, performance did not significantly differ between the contrast (78.35%; SD = 3.34%) and orientation (80.24%; SD = 3.66%) tasks (t_(16)_ = 1.796; p = 0.122). Importantly, with this larger sample we found that participant’s memory recall performance differed significantly between the 3 distractor attention conditions (**Fig. 4B**; F_(2,32)_ = 12.021, p < 0.001). Specifically, the absolute recall errors were lowest when participants ignored the distractor compared to when they attended either contrast (t_(16)_ = 3.419, p = 0.002) or orientation (t_(16)_=3.723, p < 0.001) changes in the distractor. The absolute recall error did not differ significantly between contrast and orientation attention trials (t_(16)_=1.779, p = 0.1). Thus, in this larger data set, we see the same ordinal ranking of recall errors from (1) ignore, (2) attend contrast, and (3) attend orientation that we observed in the scanner, but with more power the recall error when participants attend contrast is now significantly higher than when they ignore the distractor. In the supplemental materials (**Supp. Fig. 5**), we also document trends in the impact of attention to distractors on the classic oblique and the cardinal bias effects, as well how attention to distractors impacts attractive biases toward the distractor.

**Figure 4.**
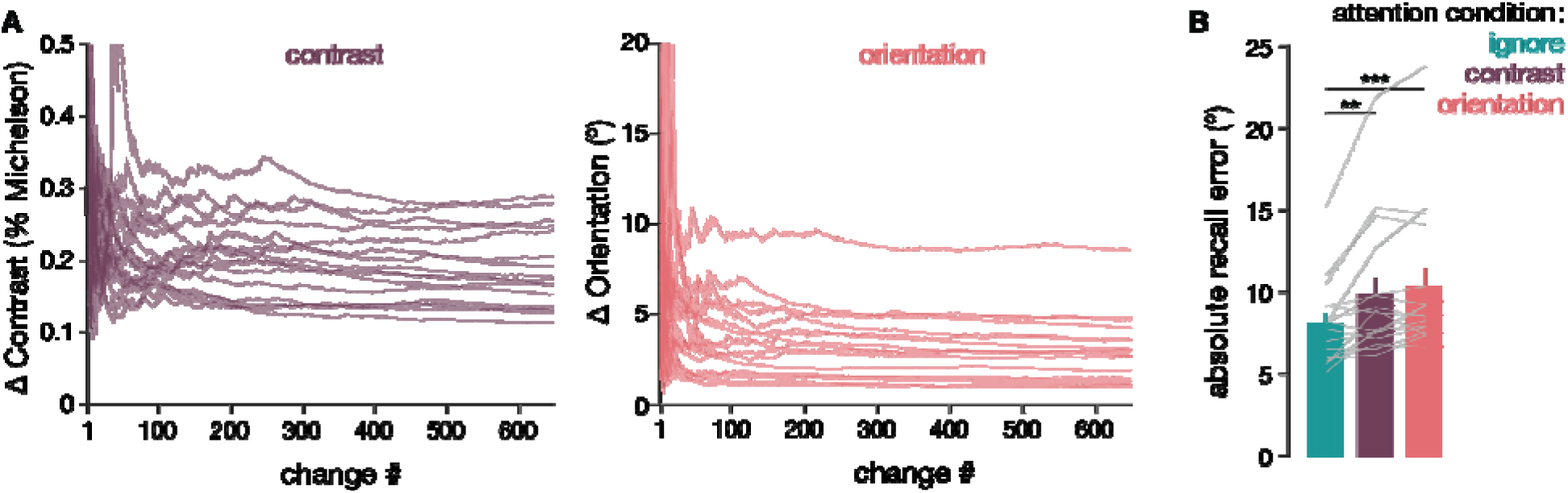
Results from the additional psychophysical experiment (**a**) Performance on the attention tas was staircased throughout the experiment. Note that a relevant change (i.e., a change in contrast when contrast was attended) could happen 2–4 times per trial, so 3 times on average. The contrast or orientation offset was adjusted with every relevant change to maintain a performance of ∼75%. While the initially noisy staircases could have impacted overall performance levels, performance did nevertheless not significantly differ between the two tasks (t_(16)_ = 1.796; p = 0.122). Lines indicate individual participants. (**b**) Recall of the memory target differed between conditions (F_(2,32)_ = 12.021, p < 0.001), and was best when participants could ignore the concurrently presented distractor grating. Bars indicate average performance, and grey lines individual participants. Asterisks indicate significant post-ho differences between conditions.

## Discussion

Holding in mind relevant sensory information for future use is a critical cognitive function, and to be useful these memories must be insulated from being overwritten by new sensory inputs. Prior studies in humans using fMRI, and in animal model systems using invasive recordings, suggest that the same general areas of early sensory cortex that support early perceptual processing are also involved in maintaining sustained memory representations (Harrison and Tong, 2009; Serences et al., 2009; Van Kerkoerle et al., 2017; Huang et al., 2024; Yiling et al., 2024). This apparent multiplexing poses a computational challenge as it is unclear how remembered information interacts with new sensory inputs, especially those that are behaviorally relevant and require prioritization via top-down attentional modulations. Here we show that memory recall was relatively spared when an ignored distractor was presented during the delay period, similar to prior reports from our lab and others (Magnussen et al., 1991; Magnussen and Greenlee, 1992; Rademaker et al., 2015, 2019). However, when new inputs were behaviorally relevant and participants engaged in a contrast or orientation detection task, there was a significant drop in both behavioral and neural markers of memory fidelity (**Fig. 1b & 2a**). Importantly, the decline in the precision of mnemonic representations was accompanied by higher fidelity representations of the distractors, suggesting that attended and remembered information directly competes in areas of early visual cortex (**Fig. 2a & 3**). These graded modulations as a function of attention to different distractors suggest a tradeoff in the ability to prioritize mnemonic information over new sensory inputs (see also: Hallenbeck et al., 2021).

The present results help reconcile prior conflicting reports that presenting distractors during a memory delay interferes with working memory representations. For example, in one experiment Bettencourt and Xu (2016) found that activity in visual cortex associated with a remembered orientation was interrupted by the presentation of faces and buildings (gazebos) during the delay period. Rademaker et al. (2019) replicated this result using similar stimuli, but also found that presenting another oriented stimulus during the delay period did not always lead to interference. Based on the present observation that directly manipulating attention reveals graded levels of distractor interference, we speculate that some classes of stimuli – such as faces – may be inherently more interesting and thus more likely to attract attention compared to simple objects comprised of a low-level visual feature like an orientation (Davies and Hoffman, 2002; Langton et al., 2008; Hodsoll et al., 2011; Morrisey et al., 2019).

This framework is also consistent with models of attentional control proposing that attended stimuli automatically gain access to working memory and thus have the potential to interact with or disrupt pre-existing working memory representations (Bundesen et al., 2005). That said, more recent work in the domain of attentional capture suggests that attended items do not always enter working memory: irrelevant distractors can attract spatial attention but be gated out of working memory whereas potentially relevant distractors attract spatial attention and are represented in working memory (Hakim et al., 2019; Maxwell et al., 2021). For instance, Hakim et al (2021) demonstrated that markers of spatial attention, such as the amplitude of alpha oscillations measured with EEG, track the position of distractors irrespective of their potential relevance. In contrast, markers of encoding into working memory, such as the contra-lateral delay activity (CDA) measured with EEG, indicate that only potentially relevant stimuli are gated into working memory and thus have the potential to induce interference. These findings are generally consistent with the present results: the “ignored” distractor in our task was presumably spatially attended – at least to some degree – yet it did not interfere with working memory because it was completely irrelevant and thus gated out of working memory. However, as soon as a distractor was both attended and task-relevant, we observed interference in both behavioral performance and in the fidelity of cortical representations.

Both selective attention and working memory are thought to be sustained by top-down feedback signals from areas of parietal and pre-frontal cortex (PFC) (Desimone and Duncan, 1995; Kastner and Ungerleider, 2000). Our observation of interference between maintaining working memory representations and attending to new sensory inputs is consistent with destructive interference occurring between competing top-down modulatory signals, perhaps due to their convergence on common feedback layers in visual cortex (Van Kerkoerle et al., 2017). Moreover, recent studies on working memory for both simple features and different classes of objects suggests that this feedback may not simply maintain a “sensory-like” representation in visual cortex (Kwak and Curtis, 2022; Xu, 2023, Duan and Curtis, 2024; Chunharas et al., 2025). Instead, the geometry of mnemonic representations in visual cortex morphs over the delay period and along the early visual hierarchy to more closely resemble the geometry of categorical representations in parietal cortex. These geometric rotations towards a more parietal-like representation are consistent with other work in non-human model systems suggesting that dynamic working memory codes may serve to insulate remembered information from sensory interference (Libby and Buschman, 2021). While Xu (2023) did not present distractors during the delay period so the functional significance of the observed rotations is not clear, Chunharas et al. (2025) did use data from a task with visual distraction during the delay. It could be beneficial if the brain (e.g., parietal cortex) applied a geometric rotation to remembered information irrespective of the visual input or attentional state dictated by the environment, as a continuous readiness to possible distractions could help ensure robustness of mnemonic contents. In the current data we see that only the representational format in parietal cortex, but not in other visual areas, is shared between memories in the absence of visual input, ignored visual input, and attended visual input. Thus, parietal cortex seems to represent working memory contents in a common format irrespective of whatever else is going on. By contrast, different continuously visible sensory stimuli – such as the sensory localizer stimuli and the distractor stimuli used in the present study – evoke different response patterns in parietal cortex. Thus, parietal cortex does not seem to unequivocally represent sensory inputs in a common format. Together, this overall pattern of memory-general patterns and sensory-specific patterns is consistent with the idea that parietal cortex employs a stable format to maintain mnemonic information in working memory.

In principle the current paradigm and dataset might be well-positioned to determine if the presence of distractors increases rotational dynamics (Degutis et al., 2024).

However, the most common approach to assessing rotational dynamics in neural codes is to learn a geometry at one timepoint in the delay period and then generalize that geometry to a later timepoint in the delay period. Critically, cross-generalization only yields interpretable results if the SNR is equated at both timepoints in the delay period because if it is not, then a failure to cross-generalize may simply reflect worse signal in one epoch. Similarly, apparent geometrical differences between conditions could also be driven by differences in SNR. In our data set, we observed a precipitous drop-off of memory decoding accuracy – and thus SNR – in the presence of attended distractors, even using models trained on a timepoint-by-timepoint basis that are capable of learning new geometries at each point in the delay period (**Supp Fig. 4**). Thus, this drop in SNR – which is our key finding of interest – unfortunately also renders cross-generalization analyses difficult to interpret with confidence.

Collectively, our findings provide an explanation to reconcile prior reports of distractor resistance and distractor interference in early visual cortex during working memory: distractors do not obligatorily interfere with working memory representations, but simultaneously attending to behaviorally relevant new inputs does lead to interference. These results - in particular the joint drop in behavioral performance and in the fidelity of working memory representations – further support a role for early visual areas in simultaneously supporting multiple cognitive functions via sustained top-down modulation signals that bias ongoing sensory processing.

## Data & Code Availability

We have made all behavioral data and all preprocessed fMRI data, from each participant and ROI, available through the Open Science Framework (OSF) at https://osf.io/2wgrv. Code to generate the experimental stimuli used during data collection, and for the analyses used to generate the figures and statistics in our paper can also be found here, with an accompanying wiki.

## Acknowledgements

This work was supported by NEI R01-EY025872 (JTS), and by the European Union’s Horizon 2020 research and innovation program under the Marie Sklodowska-Curie Grant Agreement No 743941 (RLR). No AI tools were used for this work (not for coding, not for writing, etc.) and all its flaws are entirely human made.

## Supplementary Materials

**Supplementary Table 1:** For each subject, this table holds the number of voxels in each retinotopically defined ROI (numbers shown in bigger font), and the number of voxels within each ROI are visually responsive (numbers shown smaller font in brackets).

|  | V1 | V2 | V3 | V3AB | V4 | IPS0 | IPS1-3 | LO1-2 |
| --- | --- | --- | --- | --- | --- | --- | --- | --- |
| **S01** | 2373<br>(2032) | 2033<br>(1536) | 1936<br>(1504) | 1361<br>(1001) | 461<br>(327) | 691<br>(372) | 1766<br>(292) | 896<br>(470) |
| **S02** | 2541<br>(1912) | 2349<br>(1398) | 1610<br>(997) | 1013<br>(709) | 621<br>(394) | 679<br>(166) | 1626<br>(255) | 1381<br>(551) |
| **S03** | 2271<br>(1918) | 2377<br>(1565) | 2057<br>(1296) | 1552<br>(1014) | 787<br>(387) | 1183<br>(499) | 1531<br>(568) | 914<br>(396) |
| **S04** | 2690<br>(1898) | 2636<br>(1501) | 2219<br>(1396) | 1886<br>(1066) | 958<br>(414) | 1025<br>(530) | 1696<br>(1044) | 1070<br>(309) |
| **S05** | 1565<br>(1184) | 1433<br>(938) | 1545<br>(1142) | 1153<br>(804) | 763<br>(479) | 644<br>(491) | 961<br>(635) | 503<br>(299) |
| **S06** | 2107<br>(1685) | 2158<br>(1423) | 2531<br>(1719) | 2385<br>(1421) | 737<br>(313) | 1068<br>(413) | 1389<br>(145) | 677<br>(417) |
| **S07** | 2620<br>(1569) | 2401<br>(1436) | 1993<br>(1388) | 1592<br>(910) | 882<br>(402) | 1582<br>(401) | 2854<br>(394) | 1323<br>(410) |
| **S08** | 2830<br>(1728) | 2598<br>(1628) | 1836<br>(1359) | 840<br>(686) | 474<br>(388) | 397<br>(327) | 1097<br>(883) | 888<br>(533) |

**Supplementary Figure 1:**
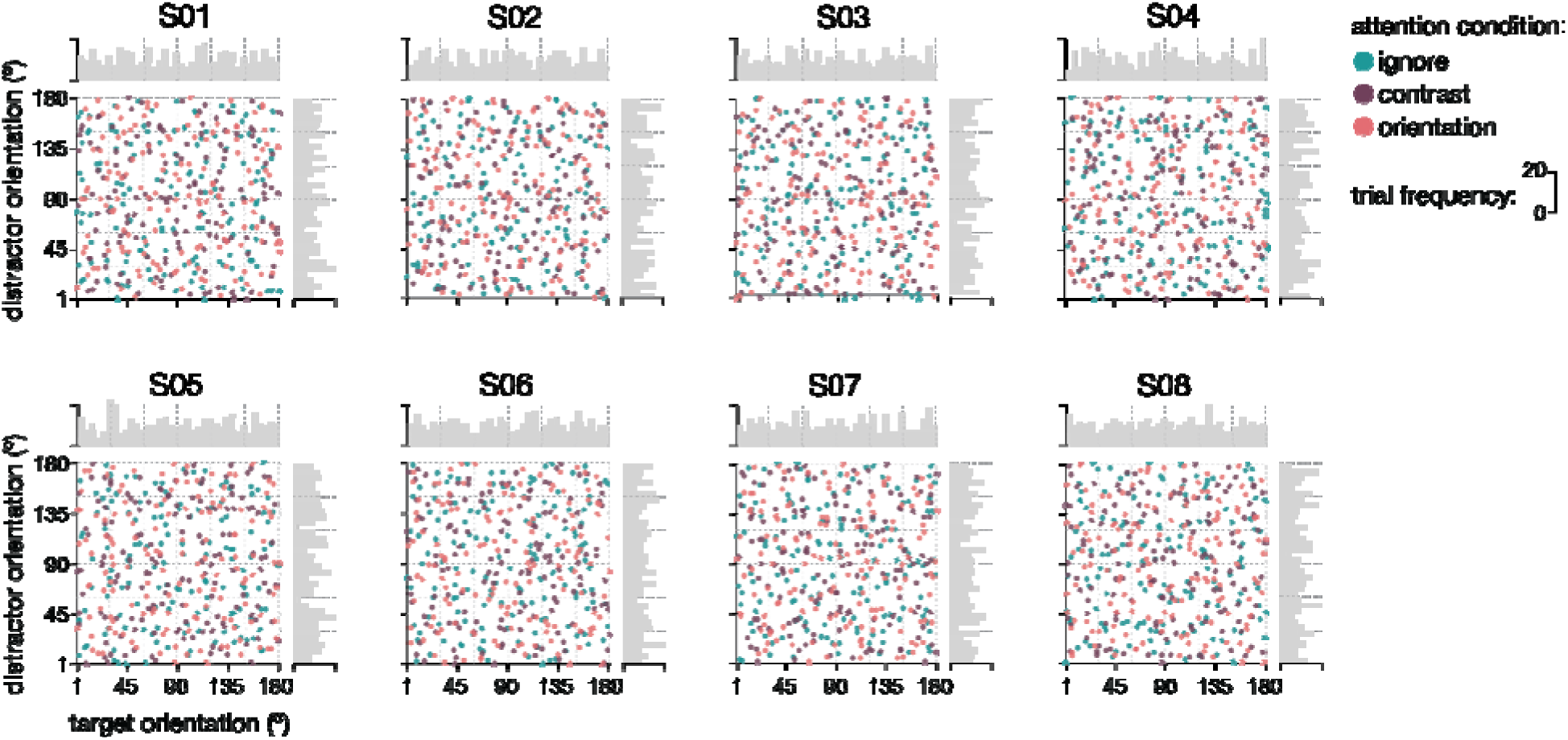
There is no systematic relationship between the orientations of the memor targets and the distractors. For each subject, here we plot the distractor orientation (y-axis) against the target orientation (x-axis) on each single trial (dots) in the main working memory experiment. Colors reflect the three different attention conditions – ignore, attend contrast, and attend orientation trials are depicted in teal, purple, and pink, respectively. To ensure relatively uniform sampling of target and distractor orientations across orientation space, both orientations were drawn pseudo-randomly from one of six orientation bins (each bin spanning 30°). The boundaries between these bins are indicated with dashed grey lines. Importantly, orientations were drawn from each bin equally often. Thus, of the 144 total trials in each condition, the target orientation was randomly drawn from the first orientation bin (1°–30°) 24 times, from the second bin (31°–60°) 24 times, and so on. Similarly, distractor orientations were randoml drawn 24 times from each bin. Importantly, draws from target and distractor orientation bins were counterbalanced, such that each bin combination (i.e., each square defined by the dashed-line grid) contains a total of 4 trials. We quantified the relationship between target and distractor orientations via circular correlation (rho), with mean correlations of –0.009 (SEM = 0.008), 0.0008 (SEM = 0.014), and 0.0104 (SEM = 0.013) for the ignore, attend contrast, and attend orientation conditions, respectively. For no subject in no condition was there ever a significant correlation between target and distractor orientations (all p-values > 0.44).

**Supplementary Figure 2:**
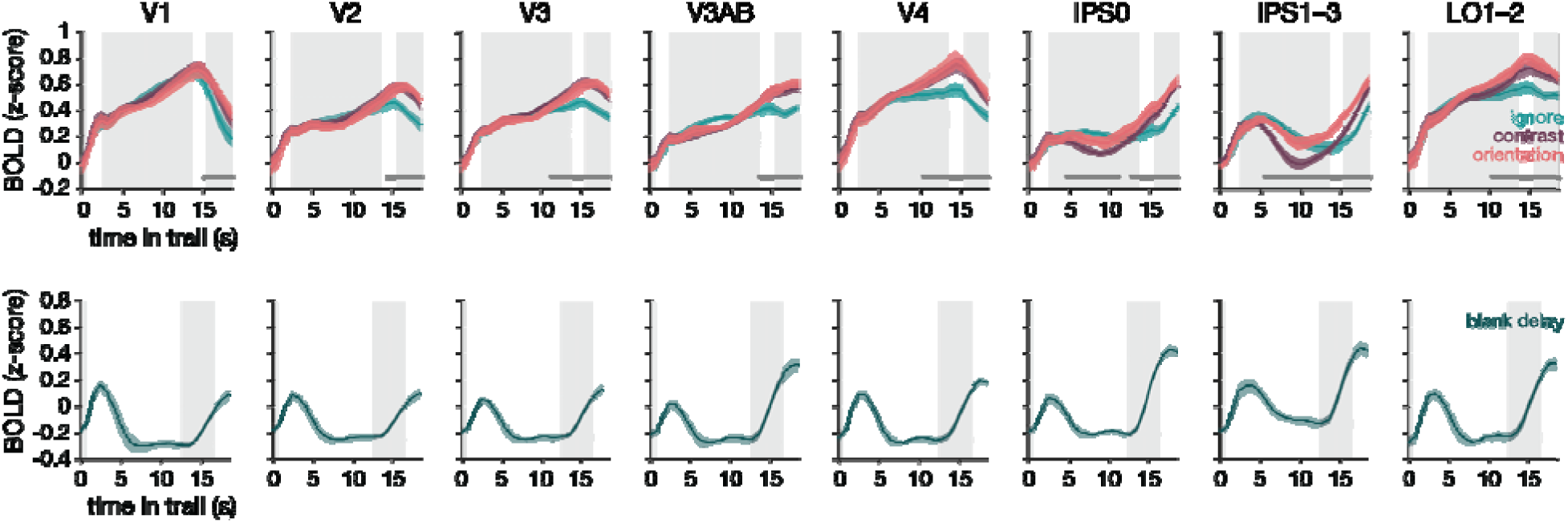
Deconvolved univariate BOLD time courses measured during the main working memory experiment (top row), and for the working memory localizer (bottom row). For the main working memory task, distractors in all 3 attention conditions effectively drove univariate responses in early visual areas (V1–V4) and LO, with overall qualitatively higher responses when attention was deployed towards the distractor. When participants attended distractor contrast (shown in purple) or distractor orientation (shown in pink), BOLD responses were generally higher then when the distractor was ignored (shown in teal), especially towards the end of the trial. In parietal regions, responses seem more transient, with the strongest response occurring with attention to distractor orientation changes (conversely, attention to contrast changes appears to result in the lowest response). Grey background-panels in each subplot indicate the time during which the memory target (0–0.5s, far left panel), the distractor (2.5–13.5s, middle panel), and the response-dial (15.5–18.5s, right most panel) were on the screen. Darker grey lines just above and parallel to the x-axis indicate clusters of consecutive TR’s during which the three attention conditions differ significantly from one another (as calculated with a cluster based permutation test, see Methods). For the memory localizer task there is only a transient response to the memory target, which goes back to baseline in the blank space between target and recall (note the absence of a grey panel during the delay of this task, and an earlier response period from 12.5–16.5s). Lines are group-averaged BOLD responses, with shaded error-bars representing + 1 within-subject SEM (N=8 independent subjects that are identical for both tasks).

**Supplementary Figure 3:**
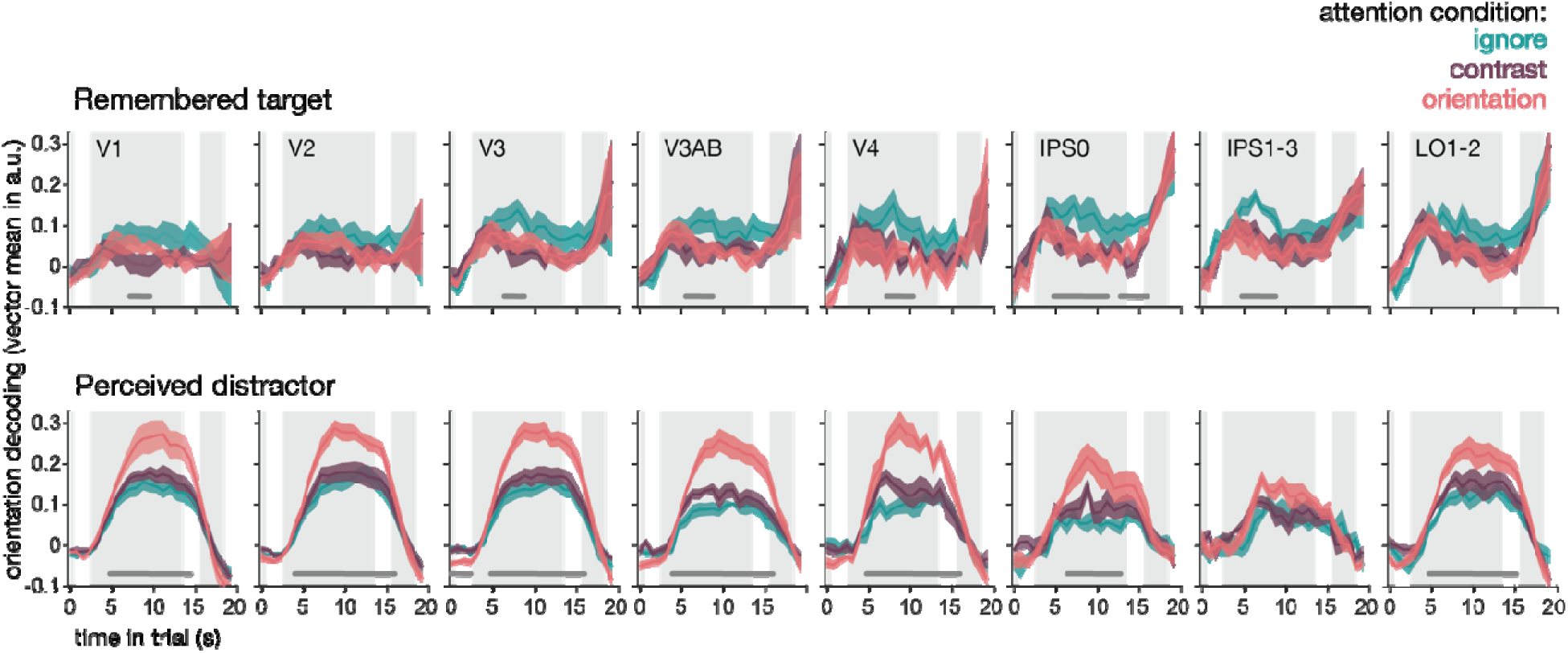
Decoding time courses for remembered and distractor orientations throughout trials of the main working memory task. The orientation held in mind during the working memory delay i better represented in visual cortex when the distractor can be ignored (teal), compared to when the distractor is attended (purple and pink, see also Fig. 2a). Here we see that these differences between attention conditions (as indicated by the dark grey lines just above and parallel to the x-axis) become apparent about 7–8 seconds into the trial. Considering the delay in BOLD response, this means differences likely emerge around the time that participants start engaging with the distractor grating (in the contrast and orientation attention conditions). The orientation of the distractor grating is best represented when its orientation is attended, intermediate when its contrast is attended, and worst when the distractor is ignored (see also Fig. 2a). Here we see that the onset of these attention condition differences happen relatively early in the trial, around 6s, which likely coincides with the very onset of the distractor grating (taking the BOLD delay into account). Grey background-panels in each subplot indicate the memor target (0–0.5s, far left panel), distractor (2.5–13.5s, middle panel), and the response (15.5–18.5s, right most panel) periods. Significance clusters indicate consecutive TR’s during which the three attention conditions differ significantly from one another (as calculated with a cluster-based permutation test, see Methods).

**Supplementary Figure 4:**
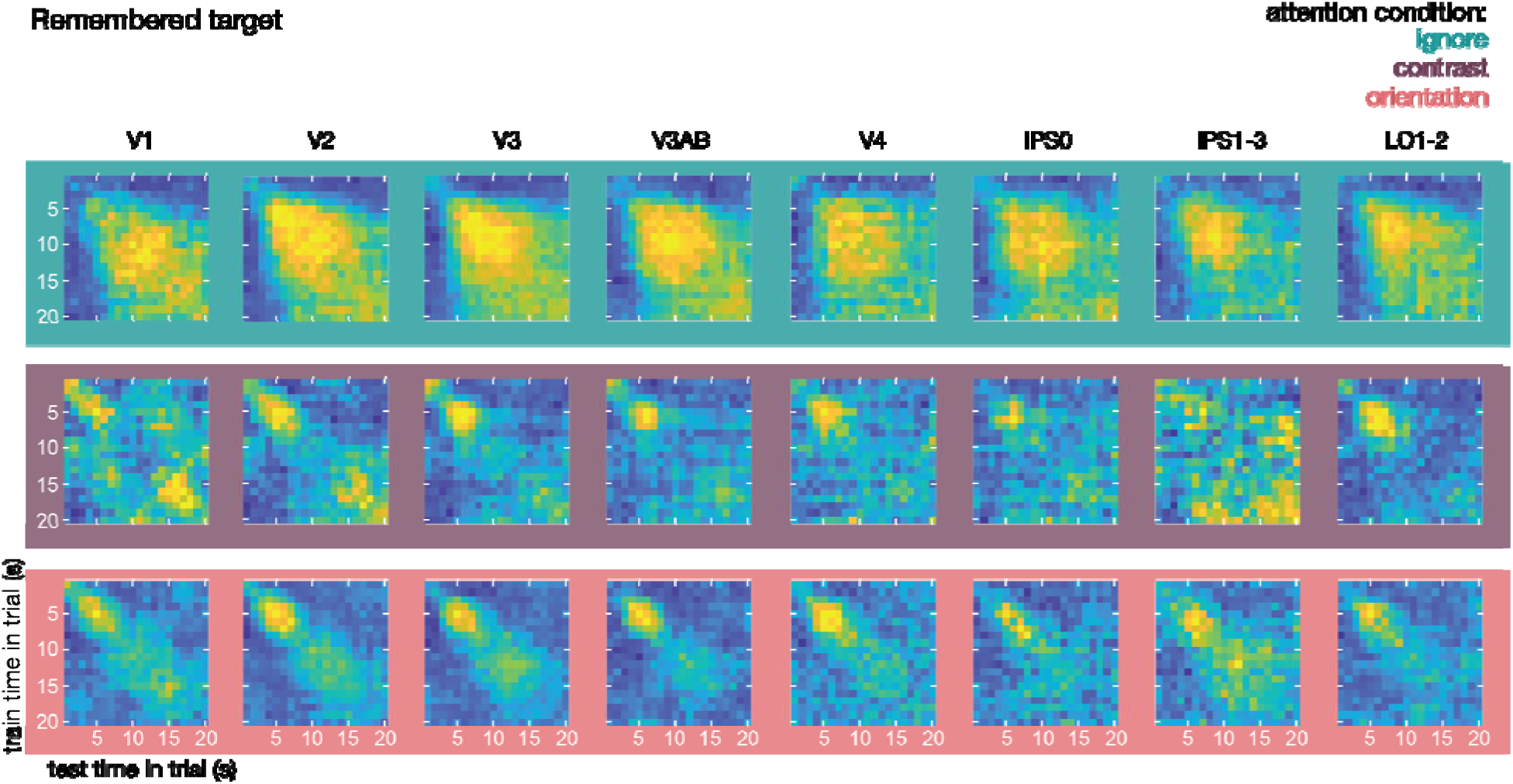
Cross-temporal decoding of the remembered orientation during the delay. Here, we train our decoder on every time point during the delay, and test it on every other timepoint. When the distractor is ignored, relatively high decoding (in yellow colors) can be seen between most timepoints during the delay, implying a sustained representation that can cross-generalize from one timepoint during the delay to many others. Under conditions where distractor contrast or orientation are attended, high decoding is more pronounced along the diagonal. However, given that decoding performance is lower in these conditions (see also Fig. 2a & Fig. 3), this could also be a matter of lower SNR when attention is diverted away from the memory stimulus.

**Supplementary Figure 5.**
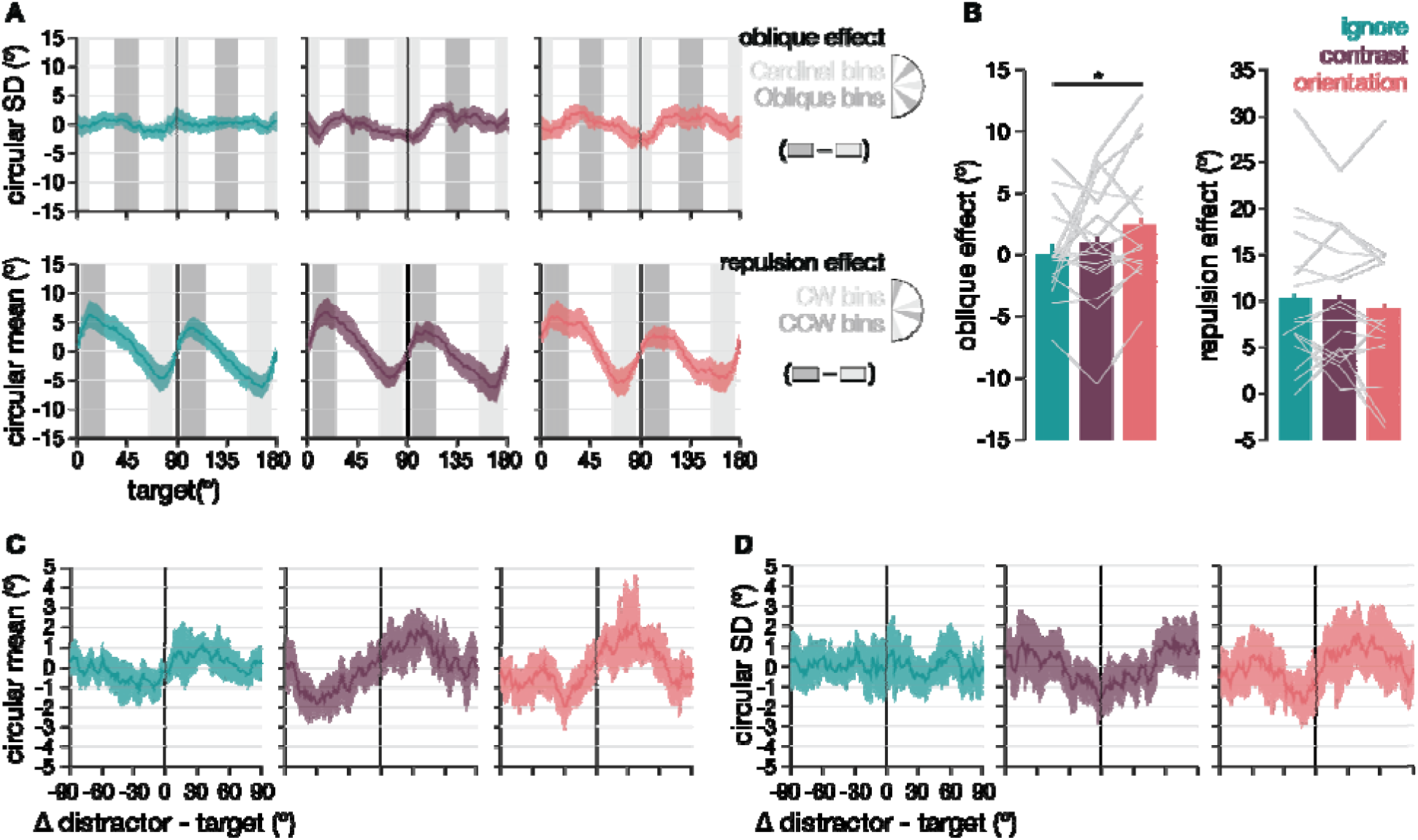
Explorative results from the additional psychophysical experiment (**a**) The top row shows the oblique effect, characterized by a smaller circular SD’s for orientations near cardinals compared to obliques. This effect appears stronger with attention to the distractor orientation (in pink). The bottom row shows the cardinal bias effect, characterized by a bias away from cardinal orientations. Because each participant performed just 216 trials per condition, we calculated the circular SD or mean at each degree + a 10° window. We normalized for between-subject differences by subtracting the average circular SD or mean for each participant before plotting and bootstrapping. Shaded areas show 95% bootstrapped CI’s. (**b**) The oblique effect magnitude was calculated by taking +10° windows around cardinal and oblique orientations, and subtracting the two such that larger numbers indicate a larger oblique effect. While a repeated measures ANOVA did not reveal a significant difference between distractor conditions (F_(2,32)_ = 2.723, p = 0.085), post-hoc tests indicated a larger oblique effect with attention to distractor orientation (t_(16)_=2.01, p = 0.046). The cardinal bias effect magnitude was calculated by taking a 20° window at 5° distance from each cardinal. The circular mean in clockwise bins was subtracted from that in counterclockwise bins, resulting in larger values for stronger bias. Cardinal bias did not differ between the three attention conditions (F_(2,32)_ = 0.939, p = 0.401). Bars indicate average performance, and grey lines individual participants. Asterisks indicate significant post-hoc differences between conditions. (**c**) The circular mean of orientation-recall responses plotted against the target-distractor offset, showing an attraction toward the distractor: When a distractor is clockwise of a target, responses are also more clockwise, and vice versa for counterclockwise responses. While this attraction appears larger with attention to the distractor, statistical evidence did not reach significance. (**d**) The circular SD of orientation-recall responses plotted against the target-distractor offset, showing that recall becomes noisier (larger SD) when the target and distractor orientations differ more. In (**c**) and (**d**), error areas indicate 95% CI’s.

## Supplemental Analysis of Psychophysical Study

There are two well-known distortions in recall for orientation stimuli, namely, the oblique effect and the cardinal bias effect (Jastrow, 1892; Lennie, 1971; Wei and Stocker, 2015, 2017; Pratte et al., 2017; Rademaker et al., 2017; Servetnik et al., 2025). The oblique effect refers to better discriminability for orientations that are closer to cardinal (i.e., 0° and 90°) compared to oblique (i.e., 45° and 135°). The cardinal bias effect refers to the bias in responses away from cardinal orientations. Together, orientation recall is more *precise* around cardinals and *biased* away from cardinals. These two effects are shown in **Supp. Fig. 5A** for all three distractor conditions (top and bottom rows are oblique and cardinal bias effects, respectively). It appears that especially the oblique effect is larger with attention to distractors. To quantify the oblique effect, we took a +10° window around cardinal orientations and oblique orientations and subtracted the circular SD of responses in those windows, such that larger values indicated a larger oblique effect. The oblique effect does not seem to differ significantly across the three attention conditions (F_(2,32)_ = 2.723, p = 0.074), although post-hoc t-tests indicate a marginally larger oblique effect magnitude with attention to distractor orientation compared to the ignore (t_(16)_=2.01, p = 0.048) and attend contrast (t_(16)_=2.03, p = 0.065) conditions (**Supp. Fig. 5B** left panel). No such differences were found for the cardinal bias effect (**Supp. Fig. 5B** right panel; F_(2,32)_ = 0.939, p = 0.41; all post-hoc p > 0.225). Although the statistical evidence is tentative, these results imply that attention to a specific visual feature (i.e., orientation) interferes with working memory for that same feature (i.e., another orientation) especially when that memory was already less precise to begin with (i.e., memory for an oblique orientation).

Recall of remembered items can be impacted by distractors shown during the delay (Magnussen et al., 1991; Magnussen and Greenlee, 1992; Rademaker et al., 2015; Bae and Luck, 2017; Lorenc et al., 2018; Kular and Serences, 2026; Ünver et al., 2026), which typically shows up as an attraction of the remembered item toward a distractor (e.g., a target is reported more clockwise in the presence of a clockwise distractor, and vice versa for a counterclockwise distractor). Additionally, responses tend to be noisier when a distractor differs more from the target (e.g., when a distractor has a 45° instead of a 5° offset relative to the target)(Rademaker et al., 2015). To examine these interactions between target and distractor, we first looked at the attraction exerted by the distractor (i.e., the circular mean of responses) as a function of target-distractor offset. In all three attention conditions we see that responses were more clockwise in the presence of clockwise distractors, and more counterclockwise in the presence of counterclockwise distractors (**Supp. Fig. 5C**). We quantified the attraction magnitude by taking the mean of clockwise responses and subtracting the mean of counterclockwise responses. We excluded a + 10° window around target-distractor offsets of Δ 0° and Δ 90°, because at these offsets there cannot *be* any attraction We did not find a significant difference in the bias toward the distractor between attention conditions (F_(2,32)_ = 2.48, p = 0.1; all post-hoc p > 0.055). Next, we looked at the precision of responses (i.e., the circular SD) as a function of target-distractor offset across all three attention conditions (**Supp. Fig. 5D**). Generally, the SD was numerically larger (i.e., precision lower) when target and distractor orientations differed more, especially when the distractor was attended. Because we had no a priori prediction about the target-distractor offset for which SD would be largest, we quantified the decline in precision by subtracting the SD when target and distractor were the same from the mean SD across all other target-distractor offsets. Quantified in this manner, the decline in precision at larger target-distractor offsets did not significantly differ between any of the three attention conditions (F_(2,32)_ = 0.944, p = 0.406). Taken together, while interactions between target and distractor exist in these psychophysical data, causing biases and impacting recall precision (**Supp. Fig. 5C and D**), attention to the distractor leads only to a small numerical amplification of these effects.

